# Bayesian species distribution models integrate presence-only and presence-absence data to predict deer distribution and relative abundance

**DOI:** 10.1101/2022.05.23.493051

**Authors:** Virginia Morera-Pujol, Philip S. Mostert, Kilian Murphy, Tim Burkitt, Barry Coad, Barry J. McMahon, Maarten Nieuwenhuis, Kevin Morelle, Alastair Ward, Simone Ciuti

## Abstract

The use of georeferenced information on the presence of a species to predict its distribution across a geographic area is one of the most common tools in management and conservation. The collection of high-quality presence-absence data through structured surveys is, however, expensive, and managers usually have more abundant low-quality presence-only data collected by citizen scientists, opportunistic observations, and culling returns for game species. Integrated Species Distribution Models (ISDMs) have been developed to make the most of the data available by combining the higher-quality, but usually less abundant and more spatially restricted presence-absence data, with the lower quality, unstructured, but usually more extensive and abundant presence-only data. Joint-likelihood ISDMs can be run in a Bayesian context using INLA (Integrated Nested Laplace Approximation) methods that allow the addition of a spatially structured random effect to account for data spatial autocorrelation. These models, however, have only been applied to simulated data so far. Here, for the first time, we apply this approach to empirical data, using presence-absence and presence-only data for the three main deer species in Ireland: red, fallow and sika deer. We collated all deer data available for the past 15 years and fitted models predicting distribution and relative abundance at a 25 km^2^ resolution across the island. Models’ predictions were associated to spatial estimate of uncertainty, allowing us to assess the quality of the model and the effect that data scarcity has on the certainty of predictions. Furthermore, we validated the three species-specific models using independent deer hunting returns. Our work clearly demonstrates the applicability of spatially-explicit ISDMs to empirical data in a Bayesian context, providing a blueprint for managers to exploit unused and seemingly unusable data that can, when modelled with the proper tools, serve to inform management and conservation policies.

## 1 Introduction

Methods to accurately predict species distributions have been central to wildlife management, conservation of endangered species, control of invasive species, and improvement of human-wildlife coexistence (Nyhus 2016, Frans et al. 2021). Species distribution models (SDMs) correlate species occurrence to variables reflecting climatic and environmental conditions, allowing us to understand spatiotemporal drivers of species occurrence in different areas or under different climatic conditions (Guisan and Zimmermann 2000). SDMs have increased in complexity since their origin, aiming to improve the predictions based on environmental variables, to account for spatial autocorrelation, and to include different data types, such as presence-only, occurrence, or presence-absence (Guisan and Thuiller 2005, Elith and Leathwick 2009, Guillera-Arroita et al. 2015).

SDMs have been developed to deal with systematically collected data with strict control for effort, methodology and spatial coverage, although these are typically expensive to collect and are thus scarce and with low spatial coverage (Hortal and Lobo 2005, Miller et al. 2019). Unstructured data, where collection effort, protocol, and exact location may not be specified, offer an alternative, more abundant even though less accurate source of information with the potential to give relevant insights about species ecology. Unstructured data may range between museum records and opportunistic citizen science observations, sometimes collected using recent advances in technology such as smartphone applications (Boyce and Corrigan 2017, Pacifici et al. 2017); in game species, unstructured data can be originated from culling returns (Nagy-Reis et al. 2021). Although unstructured datasets may be more abundant and have wider spatial and temporal coverage than structured data, their use in SDMs raises issues such as the need to carefully consider observation bias and the underestimation of local occurrence rates due to the lack of information on the observational process (Yackulic et al. 2013, Pacifici et al. 2017).

Differently from structured data, unstructured and opportunistic datasets do not include species absences and, to be used in a species distribution model, pseudo-absences need to be randomly generated in locations where the species could have been present but were not observed (Lobo et al. 2010). Although different SDM techniques have been developed to work specifically with one (e.g. presence-absences) or another (e.g. presence-only) type of data (Elith et al. 2006, Aarts et al. 2012, Isaac et al. 2019), both data types are often available for a single species, area, and time period, introducing the possibility of combining them. Two approaches have been developed to cope with this analytical challenge: data pooling and model-based data integration (or Integrated Species Distribution Models, ISDMs).

The data pooling approach combines datasets prior to entering a model, by degrading the higher quality dataset until it has a common observation process with the lower quality dataset (e.g. converting a presence-absence dataset to presence-only observations, Ahmad Suhaimi et al. 2021). Alternatively, ISDMs avoid losing data quality in the most accurate dataset by considering the two datasets as different representations of the same distribution, and thus modelling them together combining the two likelihoods (joint-likelihood approach, Pacifici et al. 2017). Additional advantages have become obvious in ISDMs: on the one hand, including an unbiased structured dataset (i.e. a presence-absence dataset) helps compensate for potential biases in presence-only datasets (Simmonds et al. 2020); on the other hand, ISDMs improve the ability to predict over a wider geographic area by combining a spatially restricted presence-absence dataset with an overlapping, but more extensive, presence-only dataset (Simmonds et al. 2020).

As datasets become increasingly complex, the challenge for SDMs is to find appropriate ways to account for the spatial structure of the observations and their intrinsic autocorrelation. Hierarchical Bayesian models allow for the inclusion of a spatially structured random effect (i.e. spatial field) that captures all the spatially explicit structures that might influence the distribution of observations (Paradinas et al. 2017, Lezama-Ochoa et al. 2020). In addition, Integrated Nested Laplace Approximation (INLA) methods have recently been implemented within the *R-INLA* package (Rue et al. 2009, Bakka et al. 2018) and the more recent development of the *inlabru* package (which provides easy access to most *R-INLA* functionality for spatially structured data, Bachl et al. 2019). INLA provides a computationally fast modelling environment for hierarchical Bayesian models where complex spatially structured random effects can be added to models for a wide variety of response variables (e.g. binomial models for presence-absence data or Poisson models for presence-only and count data, Bakka et al. 2018).

The above mentioned methods can help to model species distributions and understand their drivers, turning them into a great tool for wildlife conservation and management (Linnell and Zachos 2011). Since the latter half of the 20^th^ century, ungulate populations across Europe have shown similar expansive trends and increased local densities (Apollonio et al. 2010, Putman et al. 2011), placing them at the heart of human-wildlife coexistence research. Human-ungulate coexistence has permeated a wide variety of land-uses, among them the damage to commercial forestry plantations (Chadwick et al. 1996, Spake et al. 2020) and crops (Linnell et al. 2020); the transmission of diseases to livestock and eventually humans (Gortázar et al. 2012); and collisions with vehicles (Langbein et al. 2011). Most management plans depend on regulating the populations through hunting quotas, which requires a good assessment of population densities, locally and globally (Putman et al. 2011, Krausman and Bleich 2013, Richardson et al. 2020). However, despite the importance of having accurate estimates of population densities and distributions to inform management, survey methods are rarely coordinated or standardised, and most information comes from private stakeholders’ efforts to survey local populations (Liu and Nieuwenhuis 2014) or, at most, population estimates based on hunting returns (Apollonio et al. 2010).

Ireland provides a representative study case to apply recent advances with ISDMs to ungulate management, being home to expanding populations of native red deer (*Cervus elaphus*), and non-native fallow (*Dama dama*) and sika deer (*Cervus nippon*, Carden et al. 2011). Despite the recent population expansion of the three species (Purser et al. 2010, Liu and Nieuwenhuis 2018), Ireland lacks a national management plan for any of its deer species and, currently, management is limited to hunting permits that do not limit hunters on where (e.g. high-density hotspots), how many, and which deer (e.g. species, age and sex classes) to hunt. This is due to the lack of an empirical basis on deer distribution and relative abundance needed to set harvest quotas, maintain healthy populations and improve human-wildlife coexistence (Millspaugh et al. 2009, Williams 2011, Nagy-Reis et al. 2021). Up until now ISDMs within an INLA context had only been applied to simulated data (Simmonds et al. 2020, Ahmad Suhaimi et al. 2021); here, for the first time, we demonstrate how this approach can be applied to empirical data.

Specifically, we collated all data available on deer distribution in Ireland previously collected by several stakeholders at different spatio-temporal scales. We also collected original data using *ad hoc* web tools we created and made accessible to deer stakeholders. Our goal is to demonstrate how ISDMs can integrate structured and unstructured data to produce and validate predicted distributions for each species of deer present in Ireland, fundamental to inform science-based management practices. This study aims at demonstrating the applicability of an approach that can be adapted more broadly, and ultimately produce more accurate distributions of species that can be used for science-informed wildlife conservation and for the management of human-wildlife conflicts.

## 2 Methods

### 2.1 Studied species

There are three species of deer well distributed through Ireland, red deer, sika deer, and fallow deer. Red deer are native to Ireland (but see Carden et al. 2012), whereas fallow deer were introduced by the Anglo-Normans in the 12th century (Beglane et al. 2018) and sika deer were initially introduced for ornamental purposes in 1860s in the Wicklow mountains not far from Dublin (Powerscourt 1884).

To gather all data available on deer in Ireland and Northern Ireland (NI, UK), we contacted (1) Coillte (https://www.coillte.ie/), which provided the results of the systematic deer presence-absence surveys in part of the 440,000 ha of forests they manage in Ireland, and (2) the British Deer Society (https://bds.org.uk/), which provided survey data on the presence-absence of deer in NI. These first two datasets were the only presence-absence (PA) data available for the entire island. We collated presence-only (PO) data from (1) the British Agri-Food and Biosciences Institute (https://www.afbini.gov.uk/) which provided geotagged data on culling returns from NI. We also downloaded all observations from (2) Ireland’s National Biodiversity Database (https://biodiversityireland.ie/), a citizen science platform where users can submit deer observations, (3) iNaturalist (https://www.inaturalist.org/), an international platform with the same goal; (4) and the platform CEDaR (https://www.nmni.com/CEDaR/CEDaR-Centre-for-Environmental-Data-and-Recording.aspx) which curates all data for NI obtained from citizen science platforms and other surveys; and (5) the web survey (https://smartdeer.ie/) we developed *ad hoc* to collect PO data from Irish deer stakeholders

We obtained a total of 29,140 PA observations and 4,185 PO observations, spanning between 2007 and 2022 (the vast majority being collected in the last decade, see Table 1 for full details on the temporal resolution of data). From these, we generated three separate datasets, one for each species (red, sika, and fallow deer), to run one model for each. In addition to the PO and PA data introduced above, we gathered hunting culling returns from the National Park & Wildlife Service (NPWS, https://www.npws.ie/), responsible for issuing hunting licences. Culling returns are an alternative source of data (Milner et al. 2006, Forsyth et al. 2022), and we retained this dataset to validate the ISDMs we built by integrating PO and PA data.

**Table 1.** Summary of the presence-absence (PA, structured data) and presence-only (PO, unstructured data) datasets gathered for Ireland and Northern-Ireland (NI, UK).

|  |  |  |  |  |  |
| --- | --- | --- | --- | --- | --- |
| Data type | Source | Years | Red deer | Sika deer | Fallow deer |
| PA | BDS survey | 2016 | 920 | 920 | 920 |
|  | Coillte density surveys | 2007 - 2020 | 417 | 417 | 417 |
|  | Coillte desk surveys | 2010 - 2016 | 4 936 | 4 936 | 4 936 |
|  | **TOTAL** |  | **6 273** | **6 273** | **6 273** |
| PO | Citizen Science | 2005 - 2021 | 408 | 573 | 394 |
|  | AFBI culling returns | 2017 - 2021 | 7 | 169 | 259 |
|  | Smartdeer web survey | 2021 - 2022 | 507 | 460 | 528 |
|  | Others | 2001 - 2018 | 51 | 35 | 69 |
|  | **TOTAL** |  | **973** | **1 237** | **1 250** |

### 2.2 Data collection and pre-processing

#### 2.2.1 Presence absence (PA) data

PA data for each species were obtained from Coillte based on surveys performed in a fraction of the 6,000 properties they manage (Table 1), by asking property managers (who visit the forests they manage on a regular basis) whether deer were present and, if so, what species. Properties range in size from less than one to around 2,900 ha, and to assign the PA value to a specific location, we calculated the centroid of each property using the function *st_centroid()* from the package *sf* in R (Pebesma 2018). The survey was mainly performed in 2010 and 2013, in addition to further data collected between 2014 and 2016. Some properties were surveyed only once in the period 2010-2016, but for those that were surveyed more than once, the value for that location was considered “absence” if deer had never been detected in the property in any of the surveys, and “presence” in all other cases. In addition to these surveys, Coillte commissioned density surveys based on faecal pellet sampling in a subset of their properties between the years 2007 and 2020. Any non-zero densities in these data were considered “presences”, and all zeros were considered “absences”. These data were also summarised across years when a property had been repeatedly sampled, and counted as presence if deer had been detected in any of the samples (Table 1).

PA data for NI were obtained from a survey carried out by the British Deer Society in 2016. The survey divided the British territory in 100 km^2^ grid cells and deer presence was assigned based on public contributions, which were then reviewed and collated by experts. Since 100 km^2^ grid cells are quite large, we did not, as with the Coillte properties, calculate the centroid of each cell and assign the PA value of the cell to it. Instead, we randomly simulated positions within each cell and assigned the presence or absence value of the cell to each of them. We performed a sensitivity analysis to calculate an optimal number of positions that would capture the environmental variability within each cell (Suppl material S1), which was set to 5 random positions per grid cell. After processing, we obtained a total of 920 PA data across NI (Fig 1a).

**Figure 1:**
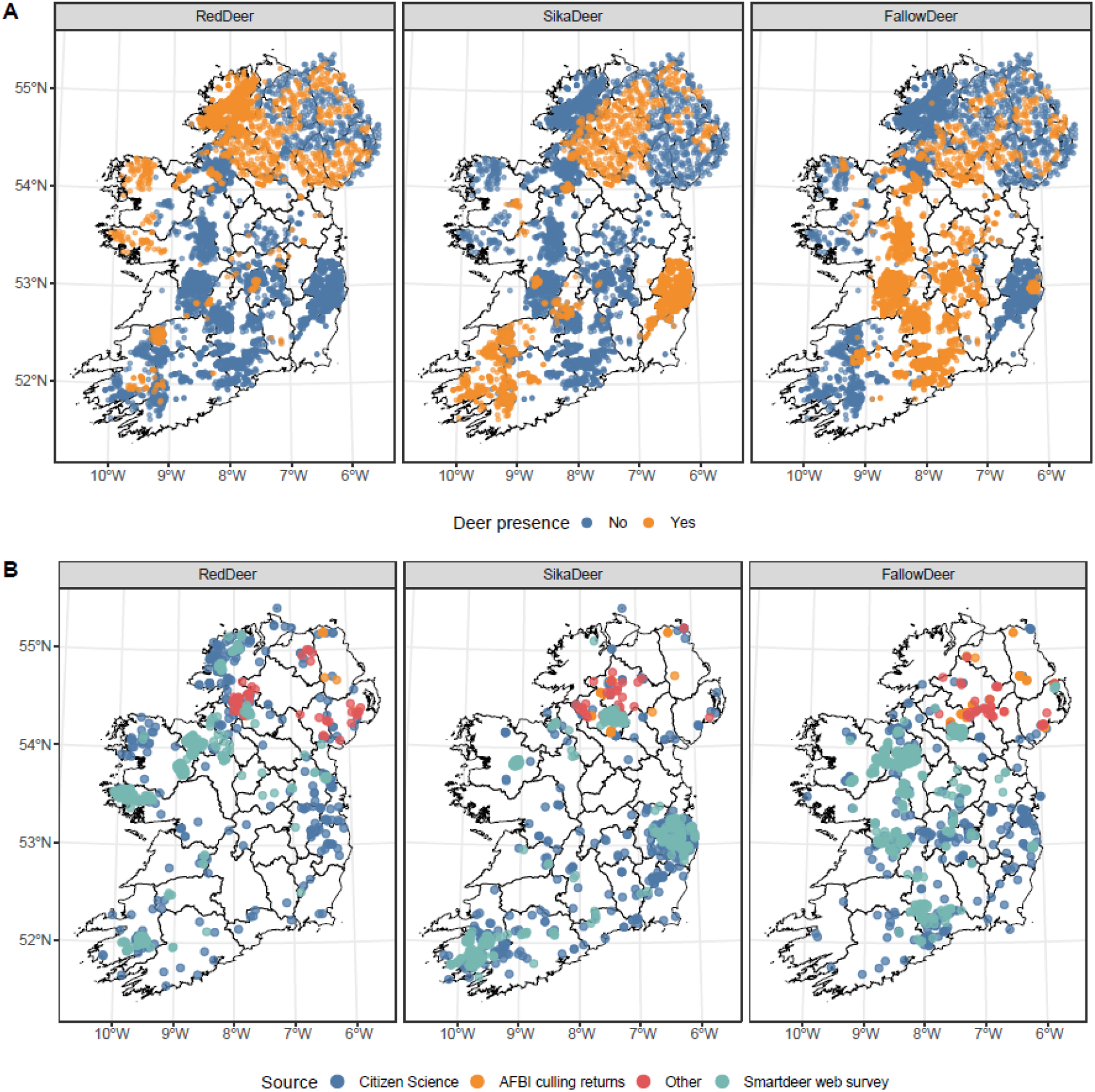
presence-absence (A) and presence-only (B) data for each deer species (see Table 1 for sample sizes and temporal resolution). Presence-absence data were provided by Coillte in Ireland and by the British Deer Society in Northern Ireland (NI, UK), while presence-only data were collated from a wide variety of sources including citizen science data, location of culled animals, and our own web tools specifically designed for deer stakeholders (https://smartdeer.ie/).

#### 2.2.2 Presence only (PO) data

PO data were collected from various sources, mainly (but not only) from citizen science initiatives. The National Biodiversity Data Centre (NBDC) is an Irish initiative that collates biodiversity data coming from different sources, from published studies to citizen contributions. From them, we obtained all contributions on the three species, a total of 1,430 records. To this, we added the 164 records of deer in Ireland downloaded from the iNaturalist site, another citizen contributed database that collects the same type of data. From the resulting dataset, we (1) removed all observations with a spatial resolution lower than 1 km^2^; (2) did a visual inspection of the data and comments, and removed all observations that were obviously incorrect (i.e. at sea or that the comment specified it was a different species); (3) filtered out all the fallow deer reported in Dublin’s enclosed city park (Phoenix Park) to avoid biases caused by the large amount of people reporting deer from the capital; and (4) filtered duplicate observations by retaining only one observation per user, location, and day. The Centre for Environmental Data and Recording (CEDaR) plays a similar role to the NBDC in Northern Ireland. They provided 872 records of deer in NI, coming from different survey, scientific, and citizen science initiatives, from which we removed all records provided with a spatial resolution lower than 1 km^2^. The location and species of 469 deer culled between 2019 and 2021 in NI were obtained from the British Agri-Food and Biosciences Institute. For the observations that did not have specific coordinates, we derived them from the location name or postcode if provided.

As part of a nationally funded initiative to improve deer monitoring in Ireland (SMARTDEER), we developed a bespoke online tool to facilitate the reporting of deer observations by the general public and all relevant stakeholders e.g. hunters, farmers, or foresters. Observations were reported in 2021 and 2022 by clicking on a map to indicate a 1 km^2^ area where deer have been observed. For each user and session, we calculated the area of the surface covered in squares, and simulated a number of positions proportional to the size of the polygon and distributed them within it to generate a number of exact positions equivalent to the area were the user had indicated an observation (details in Supp material S1). In total, the SMARTDEER tool allowed us to collect 4,078 presences across Ireland and NI (Table 1, Fig. 1b).

The data used in the models were collected between 2007 and 2022. Deer populations expanded in Ireland until 2008 (Carden et al. 2011), and according to culling return data have somewhat stabilised since then (NPWS official data). Although the range expansion of deer species would merit further investigation, here we provide for the first time an accurate modelled distribution of the three main species of deer in Ireland, and since the data are scarce, we have made use of all available data without considering the temporal trends. A continued data collection scheme will provide enough data to study population size and range changes, but this is beyond the scope of this manuscript.

### 2.3 Statistical model

To integrate the two datasets into one model for each species, we used functions from the *PointedSDMs* package (https://github.com/PhilipMostert/PointedSDMs). This package is designed to construct ISDMs from different data sources by employing the functions from the *inlabru* package (Bachl et al. 2019), within the R-INLA framework (Rue et al. 2009). Through the functions provided in the package, we constructed a joint likelihood model, with the PA data modelled as a Bernouilli distribution (Isaac et al. 2019), modelling the probability of observing an individual at each location s

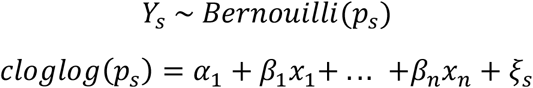

where *Y*_*s*_ is the binary response variable (PA) and *p*_*s*_ is the probability of presence. This is linked to the linear predictor by a complementary log-log link function (cloglog, Ahmad Suhaimi et al. 2021). The linear predictor is composed of a dataset-specific intercept (*α*_1_), a set of covariates (*x*_1_ *to x*_*n*_) and their coefficients (*β*_1_ *to β*_*n*_), and a dataset-specific random spatial effect (spatial field) to account for the spatial structure of the data (*ξ*_*s*_).

In turn, the PO data are modelled as a log-Gaussian Cox process with intensity function

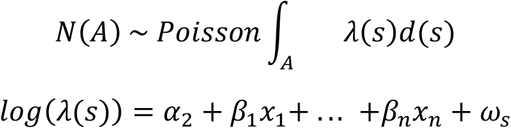

where N is the expected number of presences in the study area (A), λ is the intensity function, *α*_2_ is the dataset-specific intercept, the same vector of covariates with their effect sizes as in the PA model, and another data-specific spatial field (ω_*s*_). The use of the cloglog link in the binomial model allows its response to be interpreted on the same scale as the response of the Poisson model, which allows the sharing of parameters between likelihoods (Bowler et al. 2019). Spatial fields from both processes (PO, PA) are modelled as Gaussian random fields with Matérn covariance functions, which are approximated using a triangulation of the study area (called a mesh) through stochastic partial differential equations fitted in a Bayesian context through integrated nested Laplace approximations (Lindgren et al. 2011).

#### 2.3.1 Prior specification

The spatial fields are controlled by two hyperparameters –range and marginal variance. The range controls the smoothness of the spatial field (i.e. the distance between peaks and throughs), and the variance controls the magnitude of these peaks and throughs. In the Bayesian context in which we are fitting this model, we need to set prior values to these two hyperparameters. To do so, we use Penalised Complexity (PC) priors, a newly developed framework that allows easily interpretable and controllable priors (Simpson et al. 2014). PC priors are weakly informative (allowing the posterior of each hyperparameter to be mainly controlled by the data) and penalise model complexity by “pulling” the model towards its simplest realisation (the “base” model), which has infinite range and zero variance (i.e. a completely flat spatial field, absence of spatial structure). To set the priors, we inform the model of “how far it is allowed to deviate” from those base models using the following specifications:

- The prior on the range (ρ) is set providing the lower tail quantile *ρ*_0_ and the probability *P*(*ρ*) so that

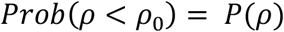

or “the probability that the true range (*ρ*) is smaller than *ρ*_0_ is *P*(*ρ*)”. For example, if we set *ρ*_0_ to be 50 and *P*(*ρ*) to be 0.05, we are telling the model that the probability of the true range of the spatial field being smaller than 50 km is 5%. In this way we are limiting the range to values between infinite (the base model) and 50, i.e. we are saying that the smallest that the range could possibly be is 50 km with a probability of 95%.
- The prior on the variance is set on the standard deviation, providing the upper tail quantile *σ*_0_ and the probability *P*(*σ*) so that

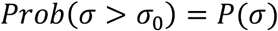

or “the probability that the true standard deviation *σ* is larger than *σ*_0_ is *P*(*σ*)”. For example, if we set *σ*_0_ to be 0.5 and *P*(*σ*) to be 0.05, the probability of the true standard deviation being larger than 0.5 is 5%, so effectively the standard deviation value is limited between 0 (the base model) and 0.5 with a 95% probability.

Priors have to be carefully specified, but there is no absolute rule for it, so the decisions that go into the prior choice are an essential part of the modelling process. In this case, we started off with a prior for the range (for both the PO and PA spatial fields, and for the three species) that was equal to the size of the triangles of the mesh (40 km). In this way, we are providing the minimum amount of information to the model, as we are setting the lower limit for the range as the limit of the resolution of the model. For the standard deviation, we started with a prior of 1 for all spatial fields, a value large enough to serve as an appropriate upper limit.

#### 2.3.2 Covariate selection

Raster environmental covariates used in the models were obtained from the Copernicus Land Monitoring Service (© European Union, Copernicus Land Monitoring Service 2018, European Environment Agency (EEA)), whereas the vector layers (roads, paths) were obtained from the Open Street Map service (OpenStreetMap contributors, 2017. Planet dump [Data file from January 2022]. https://planet.openstreetmap.org). Vector layers were transformed into distance layers (distance to roads, distance to paths) using the *distance()* function from the package *raster*, and into density layers (density of roads, paths) using the *rasterize()* function of the same package (Hijmans 2021). All raster layers were resampled to the lowest resolution available in the used covariates, resulting in a 1 km^2^ resolution. A full description of the process of covariate selection (including screening for collinearity) can be found in the supplementary material (Supp mat. S1). The covariates eventually used in the model were elevation (m), slope (degrees), tree cover (%), small woody feature density (%), distances to forest edge (m, positive distances indicate a location outside a forest, negative distances indicate a location within a forest), and human footprint index (Venter et al. 2016, 2018). All covariates were scaled by subtracting the mean and dividing by the standard deviation before entering the model (function *scale()* from the *raster* package).

#### 2.3.3 Spatial predictions

From the fitted models, we used the *predict()* function from the *inlabru* package to obtain predicted deer densities in a 25 km^2^ grid. Since models were fitted in a Bayesian context, the prediction obtained at each location is not a point value but a distribution, from which we can produce the mean and the standard deviation, thus obtaining a spatial estimate of the uncertainty of the prediction. We used the same function to obtain the prediction of the spatial effects, which can provide an indication of the spatial autocorrelation structure of each of the datasets. The model is designed on the assumption that not all individuals have been observed and although in theory the total abundance can be predicted integrating the intensity of the process over all the study area, an imperfect detection will affect the predicted total abundance. In all our models the total predicted abundances were grossly underestimated, so we decided to use the predictions in the linear scale and, rescaled from 0 to 1, use them as relative abundances instead of total abundances or densities.

### 2.4 Model validation

To validate the results of our models, we obtained culling return data from the NPWS, aggregated by county between 2008 and 2018. The data consist of the number of harvested deer of each species by county (ranging from 826 to 7,500 km^2^) and year, and the number of hunting licences issued. To consider the increase in hunting pressure affecting the number of deer harvested, we corrected each year and species data by the number of licences issued, and then aggregated the data of the past 10 years by calculating the mean. Thus, we obtained average deer harvested (corrected by number of licences) for each county. From our ISDMs, we obtained the predictions this time in the response scale, to obtain aggregated abundances by county, and then used a linear model to investigate how well our models predicted county-level culling returns, using the R^2^ score to evaluate the performance of the models.

## 3 Results

We developed one model for each species, including effects for six covariates (tree cover, density of small woody features, distance to the forest edge, slope, elevation, and human footprint index), and two spatial fields, one for the PO data and one for the PA data (Fig. S6). For red and sika deer the priors specified above for the spatial fields provided good enough posterior estimates, and we did not modify them to allow the data distribution to inform the model output. For fallow deer, a standard deviation of one proved insufficient to capture the variability of the PO spatial field, so we ran the model again with a prior value of two (Table 2).

**Table 2.**
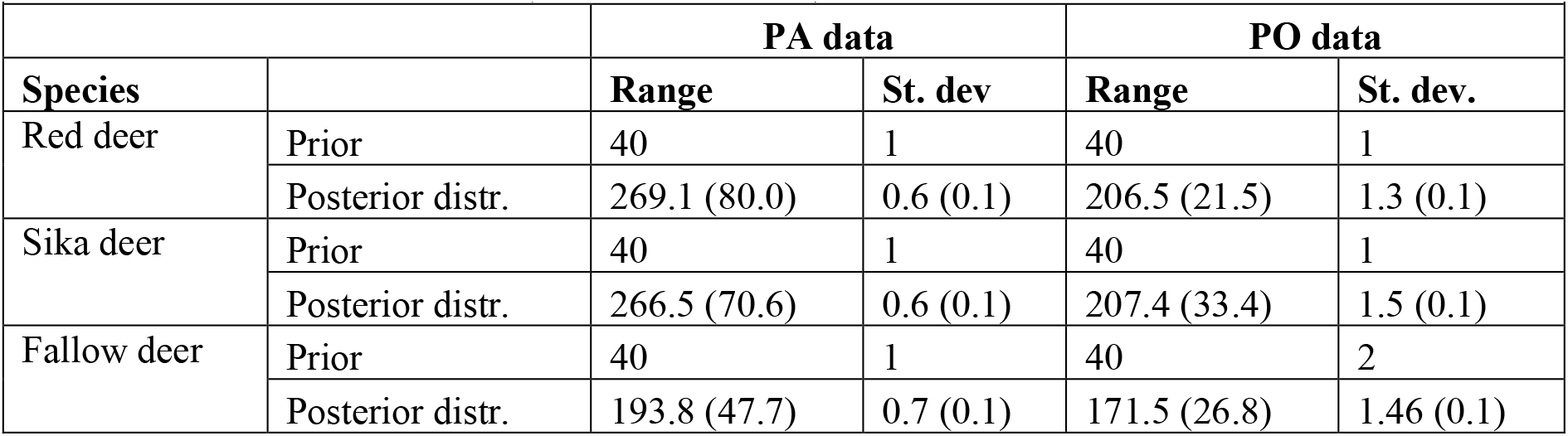
Priors’ specification and posterior distribution estimated for the spatial fields in the three species-specific models (red, sika, fallow deer) for both presence-absence (PA) and present-only (PO) data. Spatial fields are defined in our model by their range and standard deviation (St. dev). Priors are set on these parameters as point values, and posterior distributions are obtained which we have summarised here as “mean (standard deviation)”.

|  |  | PA data |  | PO data |  |
| --- | --- | --- | --- | --- | --- |
| Species |  | Range | St. dev | Range | St. dev. |
| Red deer | Prior | 40 | 1 | 40 | 1 |
|  | Posterior distr. | 269.1 (80.0) | 0.6 (0.1) | 206.5 (21.5) | 1.3 (0.1) |
| Sika deer | Prior | 40 | 1 | 40 | 1 |
|  | Posterior distr. | 266.5 (70.6) | 0.6 (0.1) | 207.4 (33.4) | 1.5 (0.1) |
| Fallow deer | Prior | 40 | 1 | 40 | 2 |
|  | Posterior distr. | 193.8 (47.7) | 0.7 (0.1) | 171.5 (26.8) | 1.46 (0.1) |

The posterior range and SD of the spatial fields showed larger ranges and smaller SD for the PA data than for the PO data, reflecting the differences in spatial structure of each dataset. PA data points are more evenly distributed throughout, while PO data points display more clustering.

The covariate effects for the three models (Fig. 2) showed that the three species had, in general, similar ecology in terms of environmental preferences, i.e. sika, red, and fallow deer were more likely to be observed within forests (negative values of distance to forest edge) with high tree cover densities. Elevation had a small but significantly negative effect on the distribution of the three species, and while slope did not have a clear effect in red and fallow deer distribution (CIs overlap zero), sika deer seemed to prefer areas with steeper slopes. The three species distributions seemed to match areas with greater human footprint, in line with the expectation that bare and unpopulated lands are less attractive to deer.

**Figure 2.**
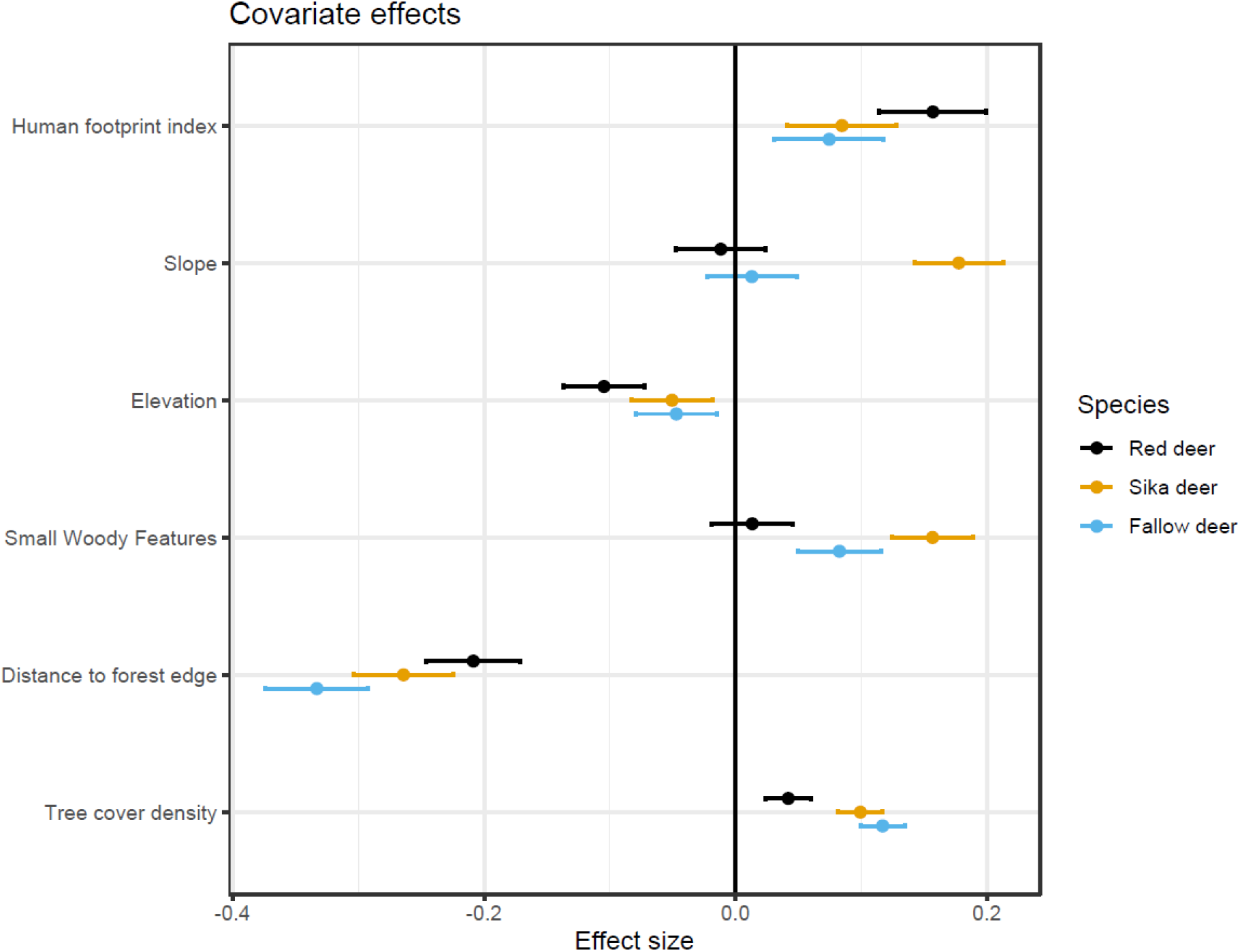
Covariate effects for each of the models for red (top, black), sika (middle, orange) and fallow (bottom, blue) deer. Circles represent the median value of the effect, while the bars represent the 95% credible intervals (CIs).

From each of the models we obtained a spatial prediction that allowed us to plot a mean prediction and its standard deviation (Fig. 3). Red deer hotspots were detected in the NW and SW of Ireland. Sika deer were present at higher relative abundances in a hotspot at the east coast, and more diffusely in the SW, overlapping with a red deer hotspot. Lastly, fallow deer are mainly distributed in the midlands. For all species, the standard deviation was larger in NI, reflecting the scarcity of PO data in that region.

**Figure 3.**
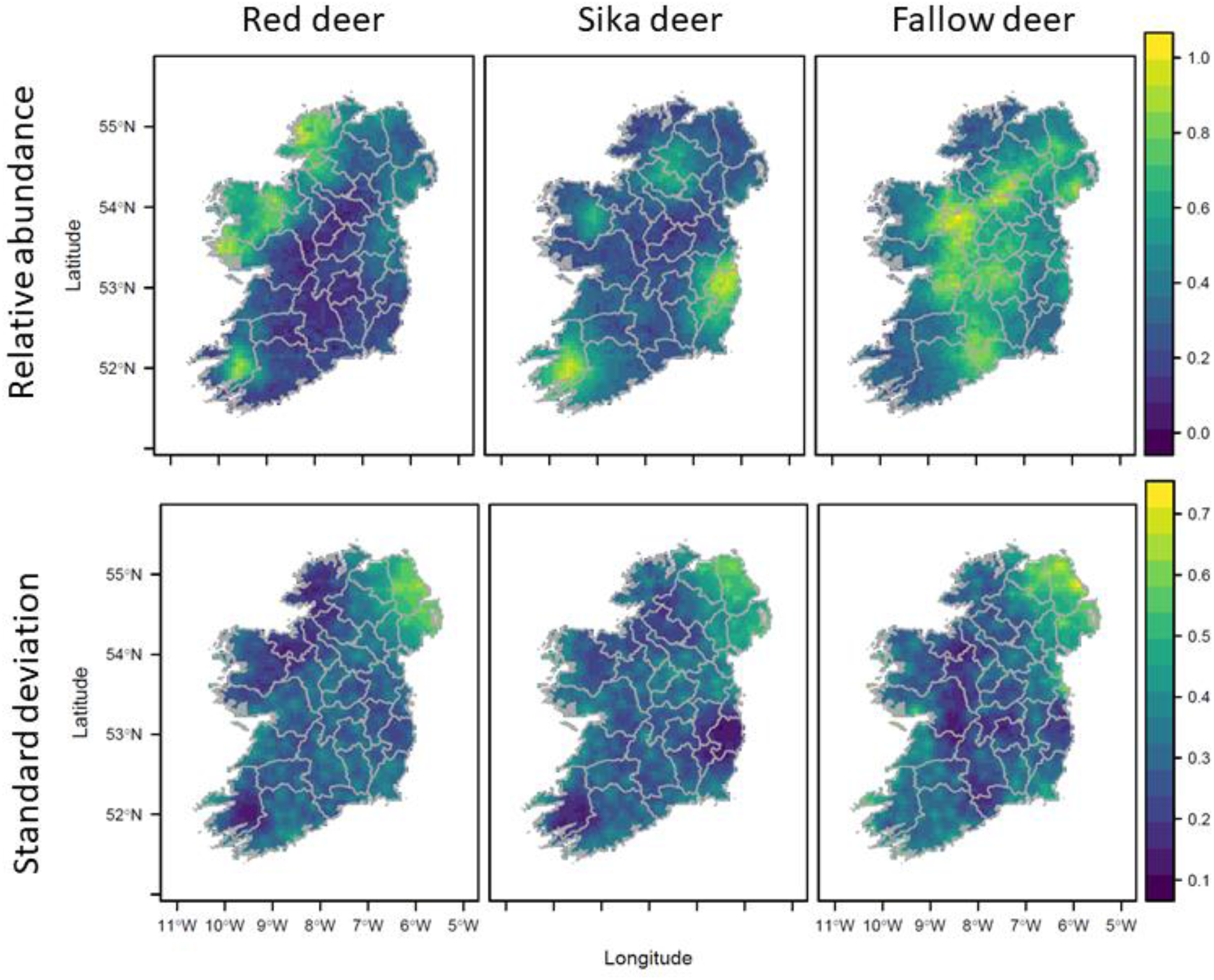
**Mean** (top) and standard deviation (bottom) of the spatial predictions for red, sika and fallow deer. The values indicate relative abundances, with 0 reflecting absence of the species and values closer to 1 representing the areas where the species is more abundant.

**Figure 4.**
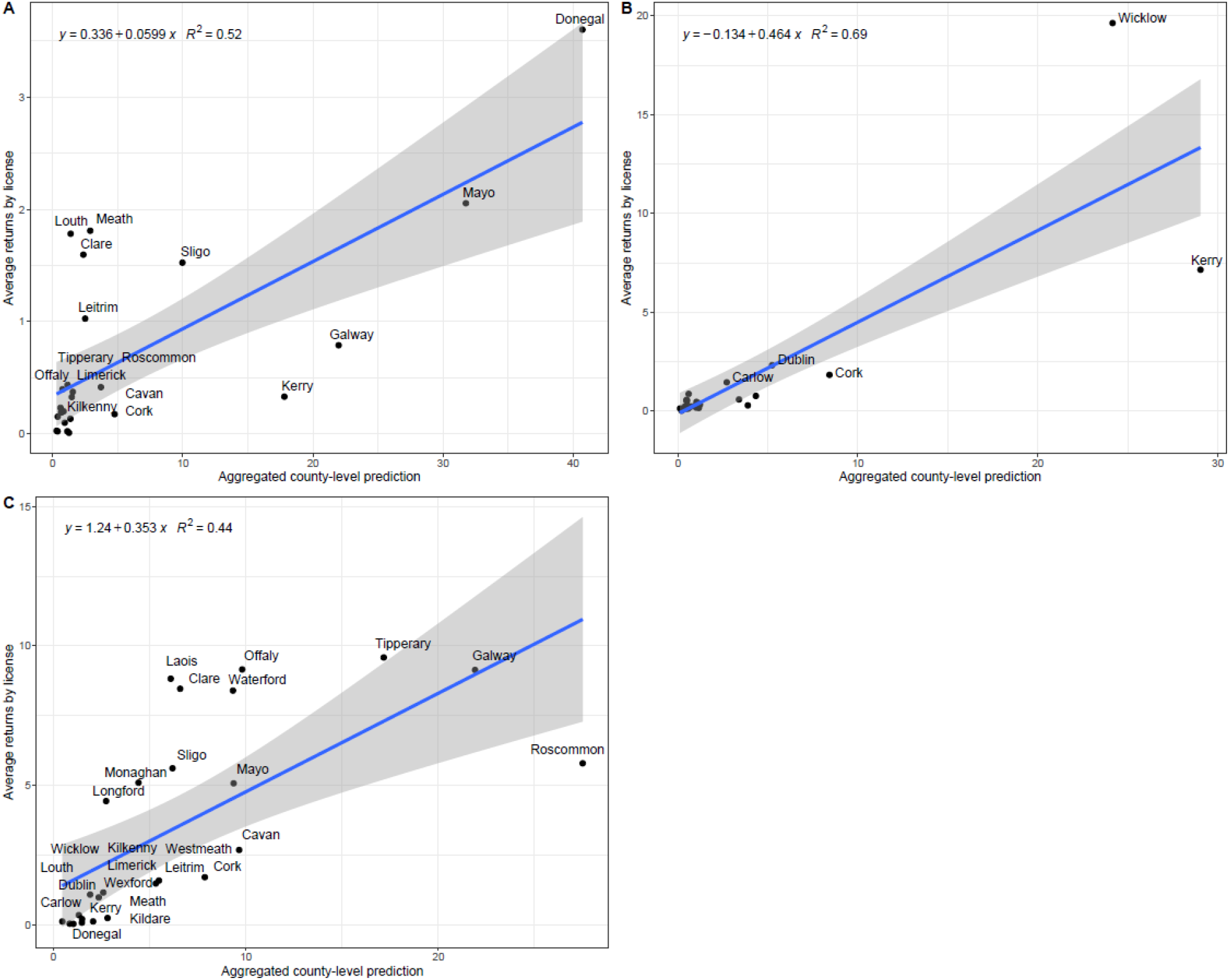
Validation plots for the ISDMs predicting red (A), sika (B) and fallow deer (C) distribution and relative abundance. Predictions of the ISDMs (x-axis, aggregated county-level abundance) are regressed against average culling returns (corrected by hunter licences at the county level, y-axis)

For the three models, ranges were larger and marginal variances smaller for the spatial fields of the PA datasets (Table 2, Fig. S1) than for those of the PO datasets, reflecting the more regular structure and thus lesser spatial autocorrelation of the dataset.

Our ISDMs predicted distribution and relative abundance across Ireland, and, when aggregated by county, these predictions were in high agreement with the independent dataset of culling returns corrected by hunters’ licences. The validation analysis showed that our models were particularly good in predicting distribution and relative abundance for sika deer (R^2^ = 0.69), followed by red (R^2^ = 0.52) and fallow deer (R^2^ = 0.44).

## 4 Discussion

### 4.1 Applicability of joint likelihood models in an INLA context to real data

Our results demonstrated the practical applications of ISDM in the INLA Bayesian context for the first time with real data, a method that so far had only been applied to simulated datasets (Simmonds et al. 2020, Ahmad Suhaimi et al. 2021). Despite the scarcity and low quality of the data, our models managed to successfully produce not only a prediction of the distribution for each species, but also to map the uncertainty. The predicted distributions displayed small standard deviations across most of the island, efficiently reflecting the regions where data are less abundant, demonstrating how fewer data relate to less certain predictions. Furthermore, we validated the predictions with an external dataset to ensure their accuracy, finding that our models performed well in predicting county-level culling returns. Thus, we provide accurate science-based relative abundance maps that integrate all previous knowledge about deer distribution in Ireland, setting a path for future data gathering initiatives with conservation and management in sight.

The separate spatial random fields for each dataset allowed us to capture the different observational processes. Although usually PA data come from organised surveys designed to avoid exhibiting any spatial structure, the PA data in our model might have exhibited some spatial structure, which would have been absorbed by the PA-specific spatial field. In the same way, PO data came from many different sources, including citizen science initiatives that would have a clear observational bias towards more populated areas or those used for recreation, but also other opportunistic observations that would have a less clearly defined observational bias. Thus, the use of a PO specific spatial field was more suited for capturing the spatial structure in that dataset than the addition of a covariate that could represent the bias, such as the human footprint index or the distance to roads (Dorazio 2014).

### 4.2 Deer distributions and relative abundances in Ireland

Our model predicted two main hotspots for red deer. The hotspot in the SW was centred around the Killarney National Park, a herd under conservation measures such as a hunting ban in the area (Carden et al. 2012). This ban is reflected in our validation plots, where our red deer model seemed to predict a larger abundance than what is reflected in the culling returns, since the culling returns of red deer in that county would be disproportionate small compared to the reality as much of their range is protected from hunting. The other hotspots to the NW coincided with areas where modern introductions of red deer have taken place in the past two centuries (Purser et al. 2010), and the diffuse populations along the eastern coast correspond to the area where the first recorded introduction of red deer into Ireland took place in 1246 (McDevitt et al. 2009).

The sika deer model showed two very clear hotspots in the E and SW of the island, and two less dense populations in the NW, reflecting the history of their introduction in Ireland (Purser et al. 2010). There was considerable overlap between the populations of red and sika deer, which could merit further study on their habitat and diet preferences to investigate the possible niche, spatial, or temporal segregation that might facilitate coexistence. From our covariate effects, sika seemed to differ in habitat preferences with red deer (non-overlapping CIs) in tree cover density and small woody feature density, where sika deer seemed to prefer denser cover than red deer, and particularly in slope, where sika seemed to prefer steeper slopes than the other two species. This difference might be reflecting some habitat or space use partitioning due to competition, but it also might be related to the fact that sika deer seem to prefer more acidic soils, which would allow them to exploit young conifer plantations (Alfredsson et al. 1998). In addition, the distribution overlap of the two species causes concerns with regards to the hybridisation between the two, which has been observed both in captivity and in the wild (Abernethy 1994) and which could be a threat to the genetic purity of the Kerry herd (Smith et al. 2014).

Fallow deer were predicted to be the most widespread species, distributed mostly over the areas from where the other two species were largely absent. This might be due to different habitat and food preferences, since fallow deer are known to be more obligated grazers than either red or sika deer (Obidziński et al. 2013), or due to competitive exclusion, but it could also be a reflection of the founder effect since fallow deer seem to have slow range expansion rates from where their populations are first established (Ward 2005). Nevertheless, since the last published distribution in 2008 (Carden et al. 2011), fallow deer distribution seems to have expanded northward, now displaying a continuous distribution from the SE coast through the midlands and the west and all the way up to the NW coast.

### 4.3 Joint likelihood models as a tool for management in data-scarce scenarios

Our predicted distributions described an island where deer of at least one species were omnipresent, with some regions where two species spatially overlap. The covariates showed that although the three species preferred areas with dense tree cover and within forests or small woody features, that did not necessarily mean that deer shy away from human presence, reflected in our models by a positive effect of human footprint index. That is, however, more reflective of Ireland’s natural habitats than of deer preferences: Ireland and NI have a large proportion of heavily modified habitat (approximately 69% of Ireland and 76% of NI are covered by farmland, (2021b, 2021a), with most of their agricultural land devoted to permanent and rough grazing grasslands, very attractive to deer (Drennan et al. 2005, O’Mara 2012), The forests, small and patchily distributed, are mostly non-native and are present within mosaics dominated by human modified habitats, making it almost impossible for deer to avoid anthropomorphised environments. This has obvious consequences for human-wildlife coexistence, since deer have more opportunities to interact heavily with human resources such as roads, commercial forestry and farms. Thus, these results constitute a starting point for management, by providing information on areas where the relative densities of the relevant deer species are higher, and where targeted actions would be most effective.

With this research, we have demonstrated the use of joint Bayesian spatial models fitted through INLA methods to obtain accurate distributions and relative abundances of species. Our models have been validated with independent data, proving their accuracy even with low quality, patchy data, which makes them a useful tool for the management and conservation of wildlife in most contexts where a data collection protocol has not been established. Our work now opens new exciting future scenarios, because the same type of model can be adapted to estimate actual abundances by including data on the number of individuals (e.g. group sizes) and sampling effort, leading ISDMs to produce even more accurate information on species abundances which are so essential for science-informed management.

## 5 Acknowledgements

We are thankful to all the agencies that provided data (complete list in methods section), and to the people that facilitated the datasets and were available to answer enquiries. Additional thanks to Sarah Keenan and Charles Harper for their contribution to the SMARTDEER project, and to Andrew McCullagh and Tony Quinn (DAFM) for their continued support throughout the development of the project.

## 7 Author contributions

VM-P: conceptualization, data curation, methodology, formal analysis, validation, visualization, writing - original draft, writing - review & editing; PSM: software, writing - review & editing; KM: conceptualization, data curation, writing - review & editing; TB: data curation, writing - review & editing; BC: data curation, writing - review & editing; BJM: conceptualization, project administration, writing - review & editing; MN: conceptualization, project administration, writing - review & editing; KM: software, writing - review & editing; AW: data curation, writing - review & editing; SC: conceptualization, data curation, supervision, funding acquisition, project administration, writing - review and editing.

