## Supplementary material for "Bayesian species distribution models integrate presence-only and presence-absence data to predict deer distribution and relative abundance"

*1 - Laboratory of Wildlife Ecology and Behaviour, School of Biology and Environmental Science, University College Dublin, Dublin, Ireland*
*2 - Department of Mathematical Sciences, NTNU Norwegian University of Science and Technology, Gløshaugen, Norway*
*3 - Deer Biologist and Management Consultant, DEER MANAGEMENT SOLUTIONS, Coolies, Muckross, Killarney, Co. Kerry V93R380*
*4 - Coillte Forest, Coillte, Dublin Road, Newtownmountkennedy, Co Wicklow, Ireland A63 DN25*
*5 - UCD School of Agriculture & Food Science, University College Dublin, Belfield, Dublin, Ireland*
*6 - UCD Forestry, School of Agriculture and Food Science, University College Dublin, Belfield, Dublin, Ireland*
*7 - Department of Migration, Max Planck Institute of Animal Behaviour, Radolfzell, Germany*
*8 - Department of Game Management and Wildlife Biology, Czech University of Life Science, Prague, Czech Republic*
*9 - School of Biology, University of Leeds, Leeds, LS2 9JT.*

#### 1 – British deer society survey: from 10 km$^2$ resolution to presence/absence points

The British Deer Society (BDS) provided presence-absence data for Northern Ireland (NI). Data were collected in a survey performed in 2016, which divided the NI territory in 100 km$^2$ sized squares, in which deer presence was assigned based on contributions by the public reviewed and collated by experts (Fig S1). The easiest way to convert these data into presence-absence data was to calculate the centroid of each square but that would mean that the environmental characteristics of the entire 100 km$^2$ square would be summarised by wherever that centroid fell (whether it was a water body, a city, or a field). Thus, we searched for a good alternative that would help better represent the environmental characteristics of each cell, so we could then relate them to the presence or absence of each species of deer. To do so, we placed random points within each cell, and extracted the values of the covariates at each point. To estimate the minimal number of points that would accurately represent the environmental diversity of each cell without artificially “oversampling” the region, we performed a sensitivity analysis.

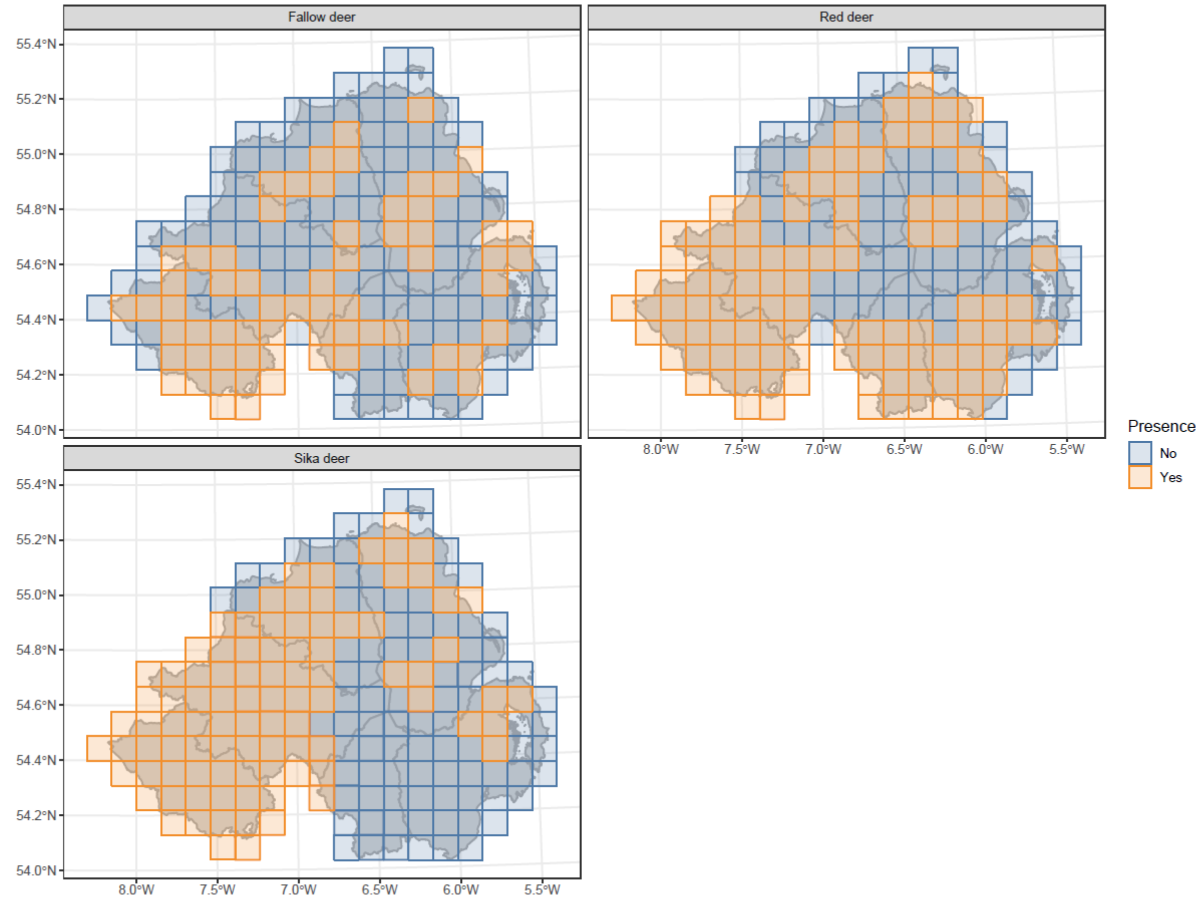


**Figure S1:** original data provided by the BDS. The maps represent the 100 km$^2$ cells in which Northern Ireland was divided, classified as “yes” where deer were observed, and as “no” where deer were not observed.

In each grid cell, we sequentially placed between 1 and 10 points randomly, and extracted the covariate values at the points. We then calculated, for each covariate and sample size, the interquartile range for each grid cell (as a measure of variability independent of sample size). We plotted those interquartile range values, and visually inspected them to search for a sample size where the interquartile range values stabilised, meaning that increasing the sample size would not increase the range of environments represented in our sample (Fig. S2). Based on those plots, we selected a value of 5 samples per grid cell – although the values for most covariates stabilised around 3 random points– to make sure we captured all available variability.

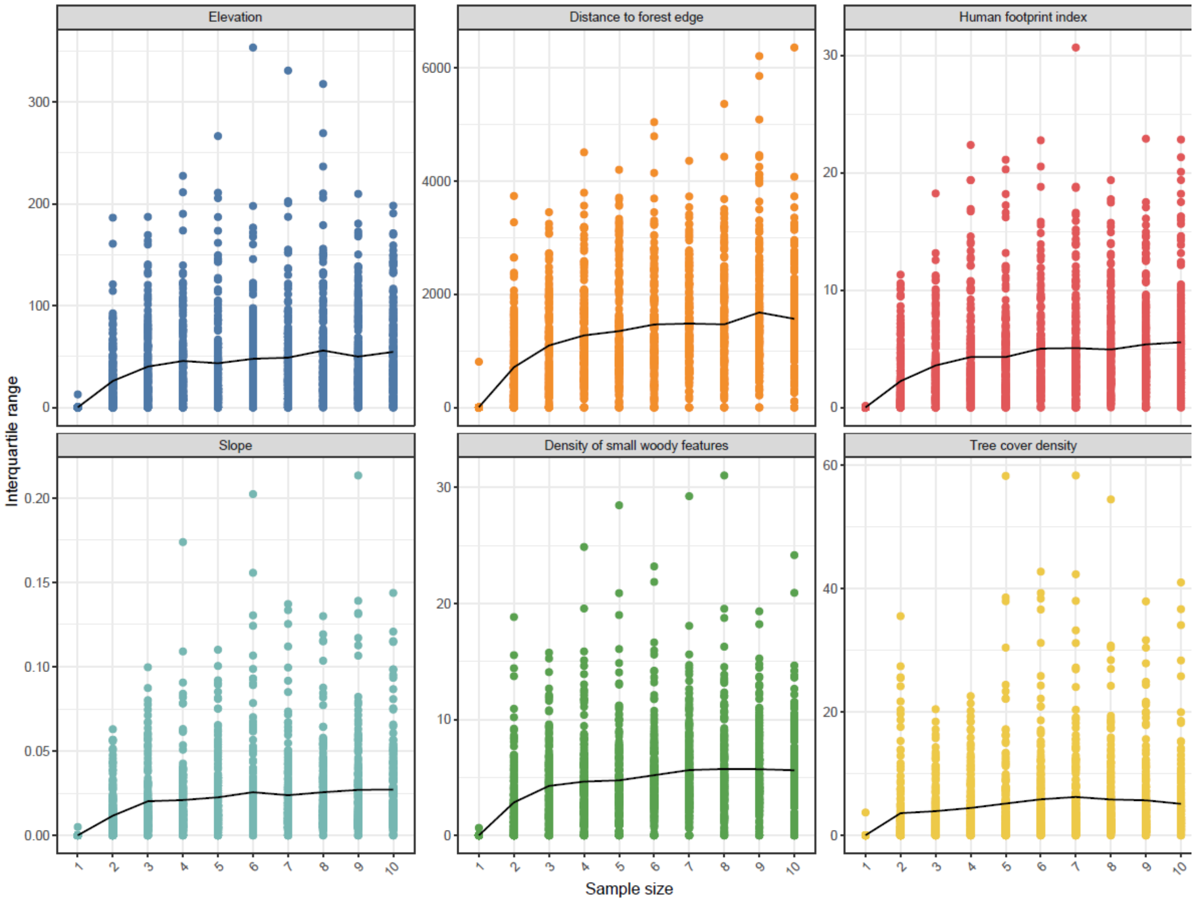


**Figure S2:** sensitivity analysis for all the covariates used as predictors in our ISDMs to decide how many random points needed to be selected within each grid cell covering the territory of Northern Ireland. For each sample size (on the x axis) the coloured circles are the interquartile ranges in each cell. The black lines represent the average interquartile range across all cells.

#### 2 – SMARTDEER web survey data: from user-selected squares to evenly distributed presence only points.

In the context of the SMARTDEER project (funded by the Irish Department of Agriculture, Fish and the Marine, DAFM), we developed a web survey where users were prompted to select, by clicking on a map, where they had seen each species of deer. Each click placed a 1 $km^2$ on the map, and thus users could indicate the entire area where deer were seen (Figure S3). The survey specifically aimed at presence-only data, and selecting one specific 1 $km^2$ cell would mean an observation of at least 1 individual of a given species. The easiest way to convert these user inputs into presence-only data is to calculate the centroid of each 1 $km^2$ square that the user has produced, which will generate as many presences as user clicks.

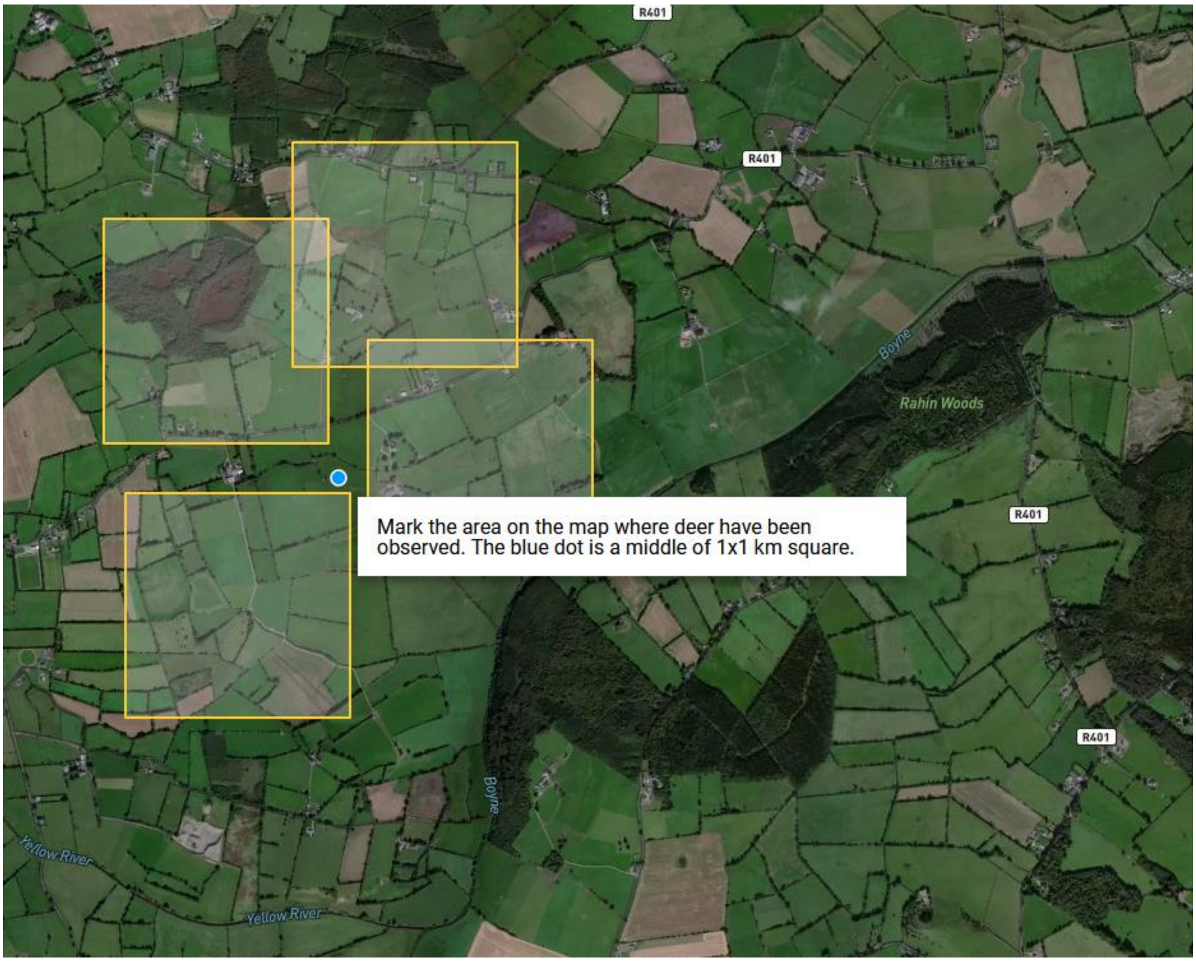


**Figure S3:** example of the typical visualisation when a user is selecting areas with deer presence (by species). The users can see the blue dot until they click, when the 1 km2 appears on the map. It is possible to click in two points in very close proximity, and thus the squares would overlap.

However, we realised that some users clicked points in very close proximity until they covered the entire area they intended to indicate (sometimes in the attempt to indicate several deer observed in the same group, or the same animal repeatedly observed in the same spot), while other users covered surveyed areas with non-overlapping squares. This generated an oversampling of some areas, since each square got a centroid, not necessarily related to a higher abundance of deer, but with the way users preferred to click in the map. To solve that issue, we dissolved all the overlapping squares generated by a single user in a single session (if there were different sessions, we assumed the user had seen the deer on separate occasions there, so we kept them separate). Once we had the surfaces composed of all the overlapping squares, we generated a number of points proportional to the area of the surface (1 point per km$^2$) and distributed them regularly across the area (Fig. S4).

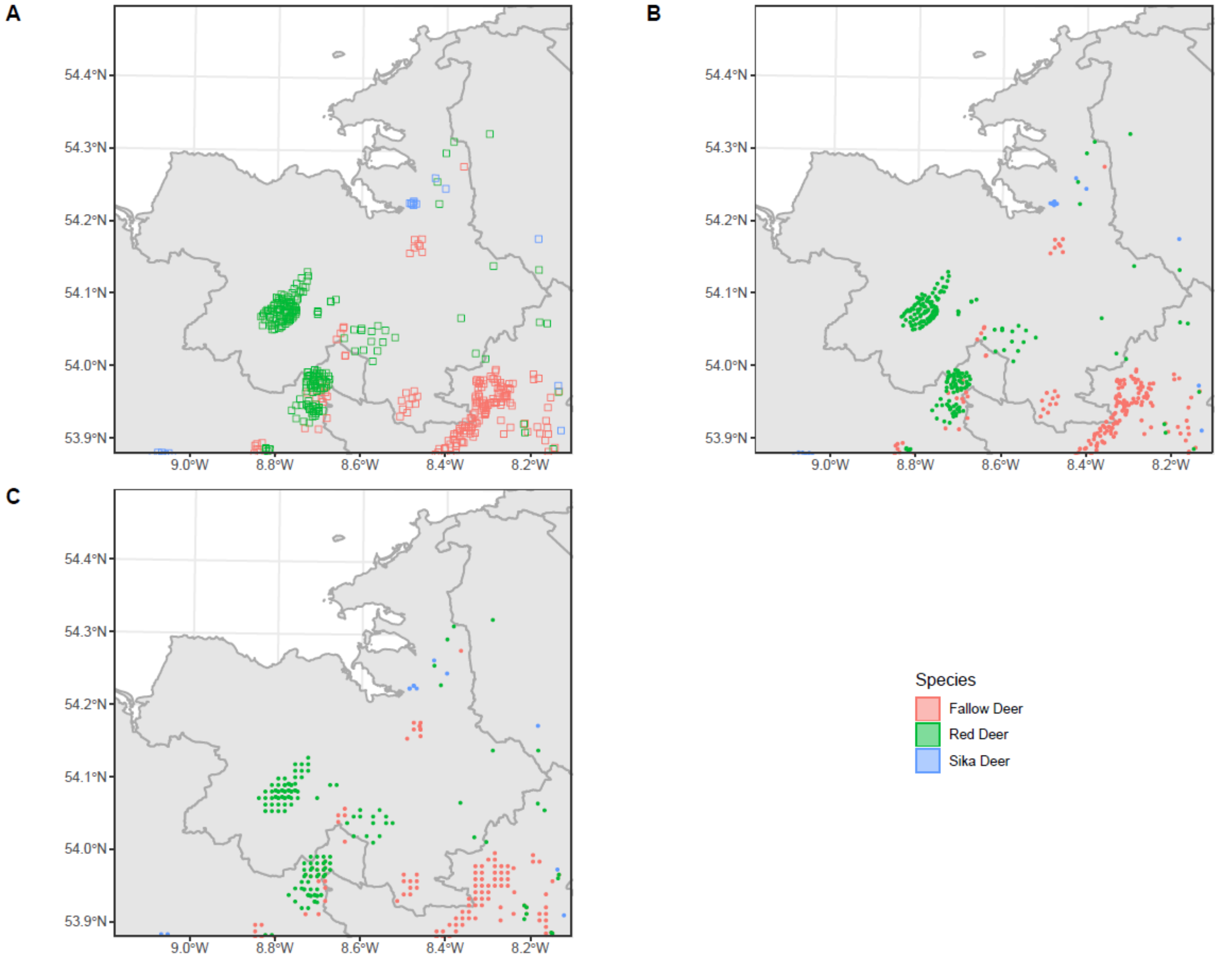


**Figure S4:** example of the resampling method used for the web survey data. Some users delimit the area where they have observed deer by clicking randomly very tight knit points until the area is covered in overlapping squares (A). If we convert the squares into points by calculating the centroid of each square, we get very close-by clustered points (B) that would overestimate the abundance of deer in that area. By contrast, if we calculate the area of the dissolved overlapping squares (by user, session, and species), we obtain artificially regular points that manage to cover the entire area where deer were observed without generating artificially high-density clusters (C). The effect of the process can be clearly seen in the area at 54.1º N and 8.6º W, where the user selected a large area as having presences of red deer by clicking many overlapping squares, but it contrasts with areas where the squares are more sparse, such as 53.95º N and 8.5ºW, where it is evident that this process does not have a strong effect in the points obtained from a user that selected presence of fallow deer by clicking non-overlapping squares.

#### 3 - Covariate selection

We considered a larger suite of covariates than those that finally entered the model, and performed a pre-selection based on correlation among them. Elevation, land use (based on the Corine map), dominant leaf type (broadleaf or conifer), tree cover density and density of small woody features were obtained from the Copernicus database (Service 2016, 2018a, b, 2019, Buchhorn et al. 2020). The human footprint index was also available online (Venter et al. 2016, 2018).

Based on the land use classification from the Corine map, we calculated distance to urban areas and distance to the forest edge, this last one taking negative values within a forest and positive outside the forest. We did not calculate inner distances within urban areas because we did not expect to find deer within them, except in urban parks (such as Phoenix Park in Dublin), and those observations were removed from the final dataset. Lastly, linear features (main, and secondary roads) were obtained from the Open Street Map (Open Street Map contributors 2017). From those, we calculated distance to paths, main, and secondary roads, and density of those features per 1 $km^2$ cell.

In a first step, we discarded all categorical layers, since unfortunately the method we were using to run the models does not allow for categorical covariates. Thus, land cover and dominant leaf type were excluded from the analysis. In a second step, we decided to simplify the covariate set. Since the human footprint index is meant to represent all human structures, we eliminated density of and distance to roads (main and secondary) as well as distance to urban areas.

With the remaining set of covariates, we calculated the Pearson correlation coefficient among them (Fig. S5). The highest correlation value was, unsurprisingly, between slope and elevation (R = 0.63), however preliminary models with and without this covariate showed that this correlation was not affecting the output, so we decided to keep it in.

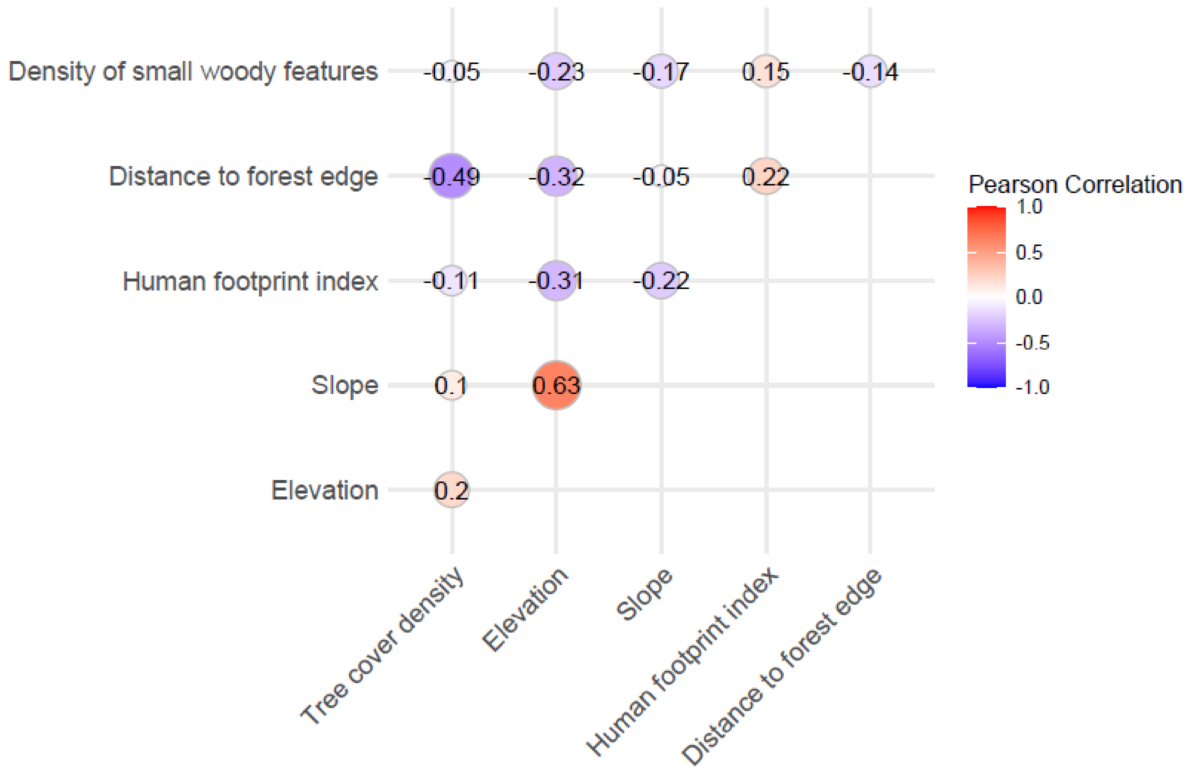


**Figure S5:** pairwise Pearson correlation coefficients between covariates. Warm colours indicate positive correlations, while cold colours indicate negative correlations. The size of the circles indicates the absolute value of the correlation. The highest correlation values are between slope and elevation (R = 0.63) and between distance to forest edge and tree cover density (R = 0.59).

### 4 – Additional figures supplementary to the main text

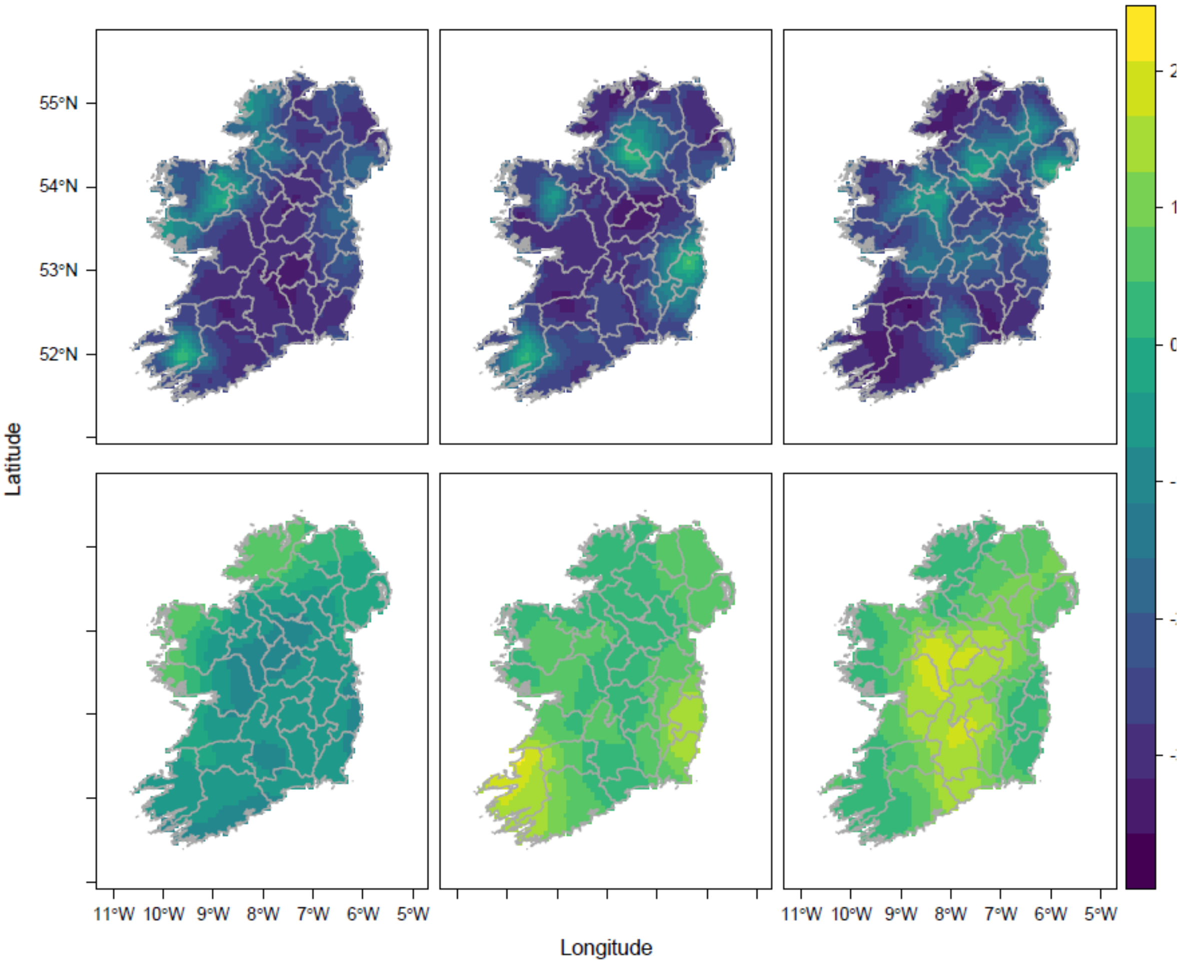


**Figure S6:** spatial fields of the models for red deer (left), sika deer (centre), and fallow deer (right). The top row displays the spatial fields for the presence-only datasets, displaying shorter ranges, and the bottom row, with larger ranges, displays the presence-absence spatial fields.

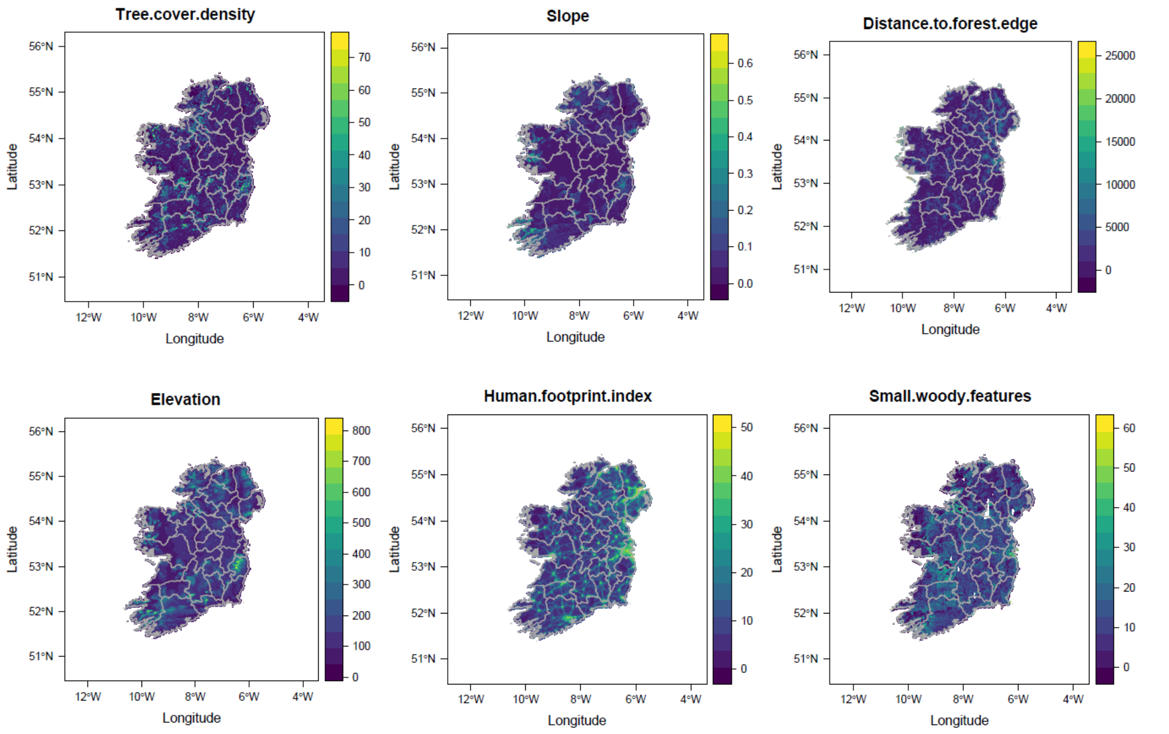


**Figure S7:** covariates entering the final models for red, sika and fallow deer. Tree cover density (in %), slope (in degrees), distance to forest edge (in m), elevation (in m), human footprint index (index value from 0 to 50) and density of small woody features (in %).

Open Street Map contributors, 2017. Planet dump. Retrieved in January 2022 from https:://planet.osm.org

Copernicus Land Monitoring Service. 2016. European Digital Elevation Model (EU-DEM), version 1.1. *European Environment Agency*

Copernicus Land Monitoring Service. 2018a. High Resolution Layer: Dominant Leaf Type (DLT) 2015. *European Environment Agency*

Copernicus Land Monitoring Service. 2018b. High Resolution Layer: Tree Cover Density (TCD) 2015. *European Environment Agency*

Copernicus Land Monitoring Service. 2019. High Resolution Layer: Small Woody Features (SWF) 2015 v. 1.2. *European Environment Agency*

Venter, O., E. W. Sanderson, A. Magrach, J. R. Allan, J. Beher, K. R. Jones, H. P. Possingham, W. F. Laurance, P. Wood, B. M. Fekete, M. A. Levy, and J. E. Watson. 2016. Sixteen Years of Change in the Global Terrestrial Human Footprint and Implications for Biodiversity Conservation. Nature Communications:12558.

Venter, O., E. W. Sanderson, A. Magrach, J. R. Allan, J. Beher, K. R. Jones, H. P. Possingham, W. F. Laurance, P. Wood, B. M. Fekete, M. A. Levy, and J. E. Watson. 2018. Last of the Wild Project, Version 3 (LWP-3): 2009 Human Footprint, 2018 Release. NASA Socioeconomic Data and Applications Center (SEDAC), Palisades, NY.
